# Cell Contractile Forces Drive Spatiotemporal Morphing in 4D Bioprinted Living Constructs

**DOI:** 10.1101/2024.09.22.613990

**Authors:** David S Cleveland, Kaelyn L. Gasvoda, Aixiang Ding, Eben Alsberg

**Affiliations:** Richard and Loan Hill Department of Biomedical Engineering, University of Illinois at Chicago, 909 S. Wolcott Ave., Chicago, IL, 60612, USA; Jesse Brown Veterans Affairs Medical Center (JBVAMC), Chicago, IL 60612, USA; Departments of Mechanical & Industrial Engineering, Orthopaedic Surgery, and Pharmacology and Regenerative Medicine, University of Illinois at Chicago, 909 S. Wolcott Ave., Chicago, IL, 60612, USA

**Keywords:** 4D bioprinting, microgel, shape morphing, biomimetics, tissue engineering

## Abstract

Current 4D materials typically rely on external stimuli such as heat or light to accomplish changes in shape, limiting the biocompatibility of these materials. Here, a composite bioink consisting of oxidized and methacrylated alginate (OMA), methacrylated gelatin (GelMA), and gelatin microspheres is developed to accomplish free-standing 4D bioprinting of cell-laden structures driven by an internal stimulus: cell-contractile forces (CCF). 4D changes in shape are directed by forming bilayer constructs consisting of one cell-free and one cell-laden layer. Human mesenchymal stem cells (hMSCs) are encapsulated to demonstrate the ability to simultaneously induce changes in shape and chondrogenic differentiation. Finally, the capability to pattern each layer of the printed constructs is exhibited to obtain complex geometric changes, including bending around two separate, non-parallel axes. Bioprinting of such 4D constructs mediated by CCF empowers the formation of more complex constructs, contributing to a greater degree of *in vitro* biomimicry of biological 4D phenomena.

## 1. Introduction

Spatiotemporal geometric transformations are essential to the development and healing processes of tissues^1^. During these processes, tissue layers undergo bending, folding, and buckling, facilitating the formation of complex geometries (**Scheme 1a-c**)^2, 3^. Cells interactions, both among themselves and with the extracellular matrix (ECM), propagate contractile forces that lead to the development of specialized tissue architectures characterized by intricate geometric shapes^4^. Cell contractile force (CCF), the mechanical force generated by the cell cytoskeletal machinery, is a driving mechanism in tissue morphogenesis steps of folding, invagination, and elongation^5^ and plays a crucial role in tissue functionalization and specialization.

**Scheme 1.**
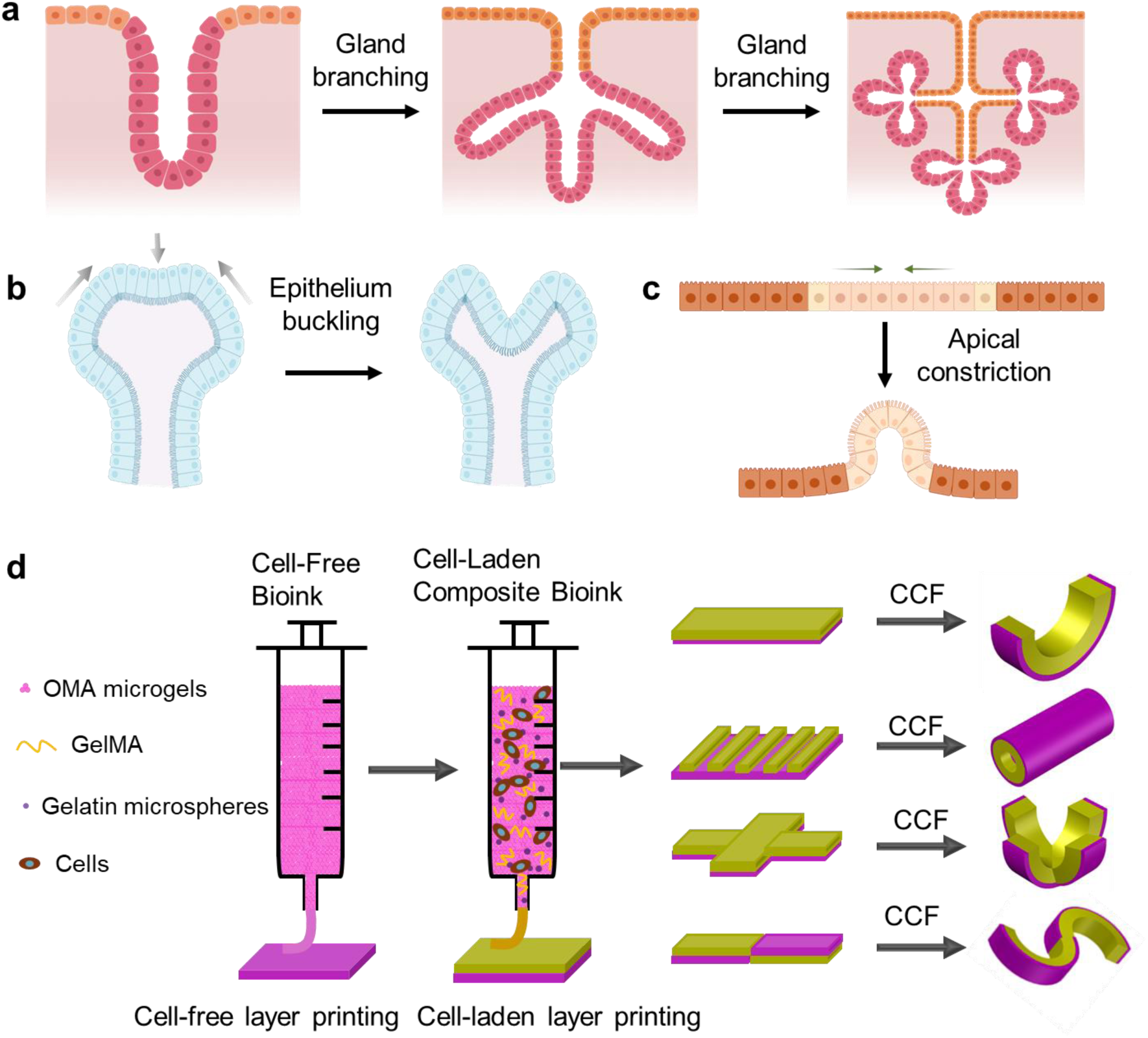
Tissue morphogenesis and 4D system design. (a-c) Typical tissue morphogenesis *in vivo*. Created with https://BioRender.com. (d) Schematic depiction of the bilayer system’s composition, fabrication methodology, and morphogenesis during culturing in media.

Recapitulating these dynamic shape changes *in vitro* holds the promise of advancing tissue engineering to a new horizon, where flexible control over geometric complexity and biomechanics can be achieved. Consequently, morphodynamical tissue engineering strategies have garnered considerable interest for their potential to produce complex, dynamic tissues that are otherwise unattainable using traditional static three-dimensional (3D) tissue engineering strategies^6, 7, 8^. Four-dimensional (4D) tissue engineering—3D tissue engineering integrated with an additional morphing component over time triggered by stimuli—has thus been developed and is rapidly evolving due to advancements in materials, instruments, and technologies^9, 10, 11^.

Current strategies in 4D tissue engineering include morphing cell condensates^12, 13, 14^ and, predominantly, morphing biomaterial matrices embedded with living cells^15, 16, 17, 18, 19^. Integrating CCF with these strategies to enable a programmed morphing process may be valuable for partially replicating tissue morphogenetic processes observed *in vivo*, thus providing effective tools for remodeling engineered tissues *in vitro*. However, morphing cell condensates based on CCF face challenges in scaling up tissue sizes and precisely controlling the morphing process. Recently, cell-only bioprinting technology has been developed to facilitate the fabrication of large cell constructs^20, 21^. When combined with a shape-morphing actuator such as a hydrogel system, 4D cell-condensate bioprinting has demonstrated effectiveness in generating large artificial tissues with complex architectures through controlled shape transformation^22^. However, this strategy relies on hydrogel swelling to elicit shape transformation rather than CCF generated within the cell condensates. Similarly, current morphing biomaterials generally depend on external energy inputs such as light, heat, electric fields, and magnetic fields to drive shape changes^23, 24, 25, 26^.

Recent advances in 4D biofabrication indicate that manipulating CCF in conjunction with biomaterials holds significant potential for generating complex living constructs. For example, seeding cells on fibronectin-coated parylene microplates can generate effective contractile forces between cells and the substrate, driving the folding of the microplates^27^. However, the cells seeded on microplates are presented in a monolayer form that exerts limited CCF, necessitating a specialized soft junction design between the nonbiodegradable microplates, which is challenging to apply in 3D tissue constructs. Alternatively, depositing loose cell clusters in the superficial region of soft Matrigel impregnated with collagen fibers via DNA Velcro technology has enabled generation of local contractile forces between cells and the surrounding collagen fibers, inducing local contraction of reconstituted tissues^28, 29^. Similarly, synthetic fibrous hydrogel assemblies composed of loose hydrogel fibers coated with cell-adhesive RGD motifs have demonstrated utility in programming shape via CCF-induced contraction^30^. While embedded cells or cell clusters within reconstituted matrices can direct global contraction and subsequent spatial tissue remodeling, these systems require ultra-weak biomaterials to allow the relatively weak CCF to exert contraction functions. Consequently, engineering initial matrix shapes is highly challenging due to the inherent fragility of the biomaterials, thereby limiting the production of final complex living constructs.

Considering the challenges associated with current engineering systems and the critical role of tissue morphogenesis in tissue development, we present the development of a CCF-driven shape-morphing living system composed of uniformly embedded living cells within a robust microgel matrix. **Scheme 1d** illustrates this system as a bilayer system consisting of a layer of photocured oxidized methacrylate alginate (OMA) microgel and a layer of photocured OMA microgel/gelatin methacrylate (GelMA) loaded with gelatin microspheres and living cells. Key features that distinguish this system from previously reported ones include: (1) the microgel materials exhibit shear-thinning and rapid self-healing properties, allowing for extrusion printing to produce stable, free-standing constructs, (2) cells are evenly distributed throughout the biomaterials, capable of generating CCF collectively to induce global construct morphing efficiently, and (3) the ability to localize CCF to induce complex tissue morphing in a precise and controllable manner. Using this system, we demonstrated advanced shape programming via CCF generated within the living constructs post-printing. Additionally, we conducted proof-of-concept 4D tissue regeneration studies, successfully demonstrating the formation of bone-like and cartilage-like tissues with preprogrammed curvatures.

## 2. Results

For 4D shape transformations to occur via CCF, the material system needs to have low mechanical properties for the cells to drive the geometric changes^31, 32^ yet durable to maintain stability after printing. To accomplish this, a system composed of OMA, GelMA, and uncrosslinked gelatin microspheres was developed. These polymers were combined to form a bioink that can be 3D printed with high fidelity while establishing a “mechanically soft” microenvironment capable of being deformed by cellular forces. To achieve printable bioinks, the OMA was processed into a jammed state which demonstrated shear-thinning and rapid self-healing^19, 22^. The GelMA was added to the polymer composition to enhance cell-ECM interactions^33, 34^, and the gelatin microspheres were incorporated to make macropores within the scaffold to: i) weaken the scaffold after they liquify, which benefits shape morphing by CCF and ii) create pores for increasing nutrient diffusion and enabling increased cell proliferation and migration, which may in turn increase the number and distribution of cell contributing to deforming the scaffold (**Fig. 1a**).

**Fig. 1.**
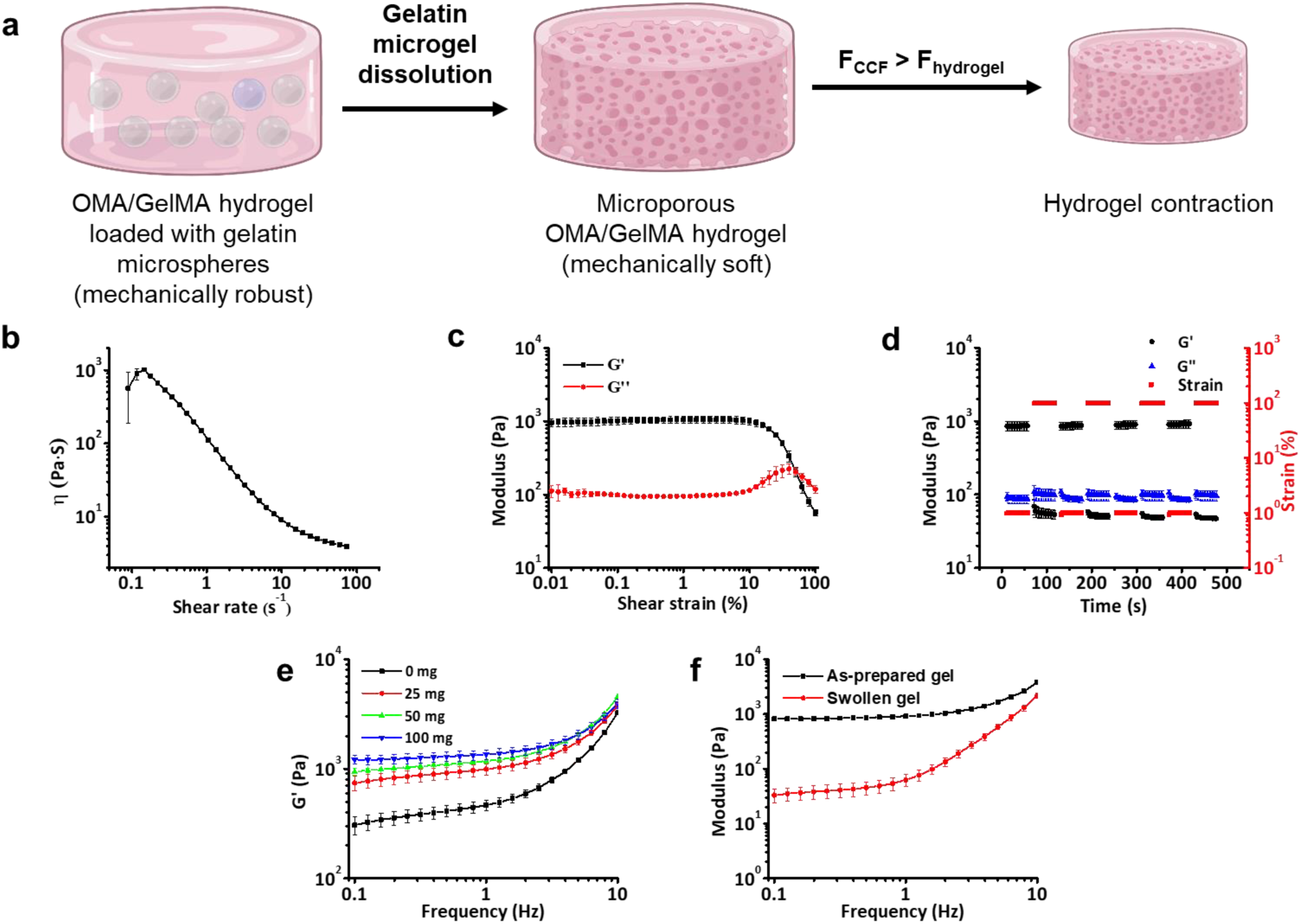
Rheology of composite OMA/GelMA bioink for 3D printing. (a) Schematic illustration of microporous matrix formation upon gelatin microsphere dissolution, which weakens the hydrogel and facilitates CCF-mediated hydrogel contraction. Created with https://BioRender.com. Composite bioink shear thinning properties, where (b) viscosity decreased as shear rate increases, and (c) the storage modulus was greater than the loss modulus at low shear strains and less than the loss modulus at high shear strains using 50 mg/mL gelatin microspheres mixed with OMA/GelMA composite bioinks. (d) OMA/GelMA composite bioink exhibited self-healing properties, with the storage and loss moduli maintaining their initial values after multiple oscillations of strain with 50 mg/mL gelatin microspheres. (e) Storage modulus (G’) increased with addition of gelatin microspheres at concentrations of 25, 50, and 100 mg/mL to the OMA/GelMA bionks. (f) Frequency sweep rheology demonstrating that after one day of culture, the 50 mg/mL of gelatin microspheres liquified in the construct composed of photocrosslinked OMA/GelMA hydrogel, causing the G’ to decrease dramatically.

To be used for free-standing 3D printing, bioinks must exhibit shear-thinning and rapid self-healing properties^20^. The viscosity of the composite microenvironment decreased dramatically as shear rate increased, confirming shear-thinning behavior (**Fig. 1b**). Additionally, the storage (G’) and loss (G’’) moduli of the bioink crossed over each other as shear strain increased (**Fig. 1c**). At low shear strains (i.e., when the bioink was at rest), G’ was greater than G’’, indicating that the bioink behavior was solid-like. However, at high shear strains (i.e., when the bioink was flowing through the needle), G’ was less than G’’, indicating that the bioink behavior became more liquid-like. Along with these shear-thinning properties, an oscilliatory strain test revealed the self-healing quality of the bioink (**Fig. 1d**). As strain oscillates between 1% and 100%, the moduli oscillated between the same values, indicating that the bioink is able to self-heal and consistently respond to reverses in strain application even after multiple exposures. Taken together, these shear-thinning and self-healing behaviors imparted the composite bioink with exceptional printability. However, since the purpose of this bioink is to aid in CCF-mediated scaffold shape changes, these rheological properties were rendered moot unless the crosslinked bioink was also soft enough to be deformed by cellular forces (G’ < 200 Pa)^35, 36, 37^. Therefore, G’ was measured at frequencies less than 10 Hz for variations of the bioink containing different concentrations of gelatin microspheres (**Fig. 1e**). The G’ of the bioink was observed to substantially increase (, whereas G’’ showed a moderate increase with increasing gelatin microsphere concentrations (**Fig. S1a**). This enhancement in the bioink’s mechanical properties underscores the beneficial impact of gelatin microspheres, promoting greater construct stability post-printing. However, since the gelatin microspheres were uncrosslinked and liquefied when cultured at 37 °C, the G’ of the photocrosslinked composite hydrogel in its swollen state after one day of culture at 37 °C was significantly reduced (**Fig. 1f**), which may facilitate cellular forces to deform the hydrogel matrix. Nevertheless, the composite hydrogel miantained its mechanical integrity, as evidenced by the larger G’ relative to G’’ (**Fig. S1b**).

To investigate the extent to which cellular forces can deform composite matrices of varied mechanics, cell-laden OMA/GelMA bioinks with various concentrations of gelatin microspheres were printed into 10 mm × 10 mm × 0.6 mm squares, crosslinked with UV light, and cultured for 14 days. Photographs of each sample (*N* = 4) were obtained each day using a dissection microscope. Representative photomicrographs for each condition at the day 1, 3, and 14 timepoints showed relative cell-mediated shrinkage of the constructs over time (**Fig. 2a**). The shrinkage of each cell-laden hydrogel was quantified by measuring the area of each square, revealing the trend that increased gelatin microsphere concentration resulted in reduced area at day 14 (**Fig. 2b**). In other words, significant contraction of the hydrogel matrix was observed at higher concentrations of gelatin microspheres. The gelatin microspheres liquefied at the beginning of culture, leaving microscale pores within the printed constructs (**Fig. S2**). The aforementioned findings were likely due to increasing concentration of micropshere producing pores reducing the mechanical resistance of the constructs to deformation by cellular forces, resulting in a noticeable increase in construct contraction. Cells in constructs in all conditions exhibited predominantly alive cells at days 1, 7, and 14 (**Fig. S3**), demonstrating high cytocompatibility of the system. While the gelatin microsphere concentration of 100 mg/mL condition conferred the greatest amount of contraction, it also posed challenges in generating reproducible, high-fidelity prints. Therefore, we determined that a gelatin microsphere concentration of 50 mg/mL was the optimal concentration to ensure macroscopic changes in shape while maintaining hydrogel stability after microsphere liquefaction.

**Fig. 2.**
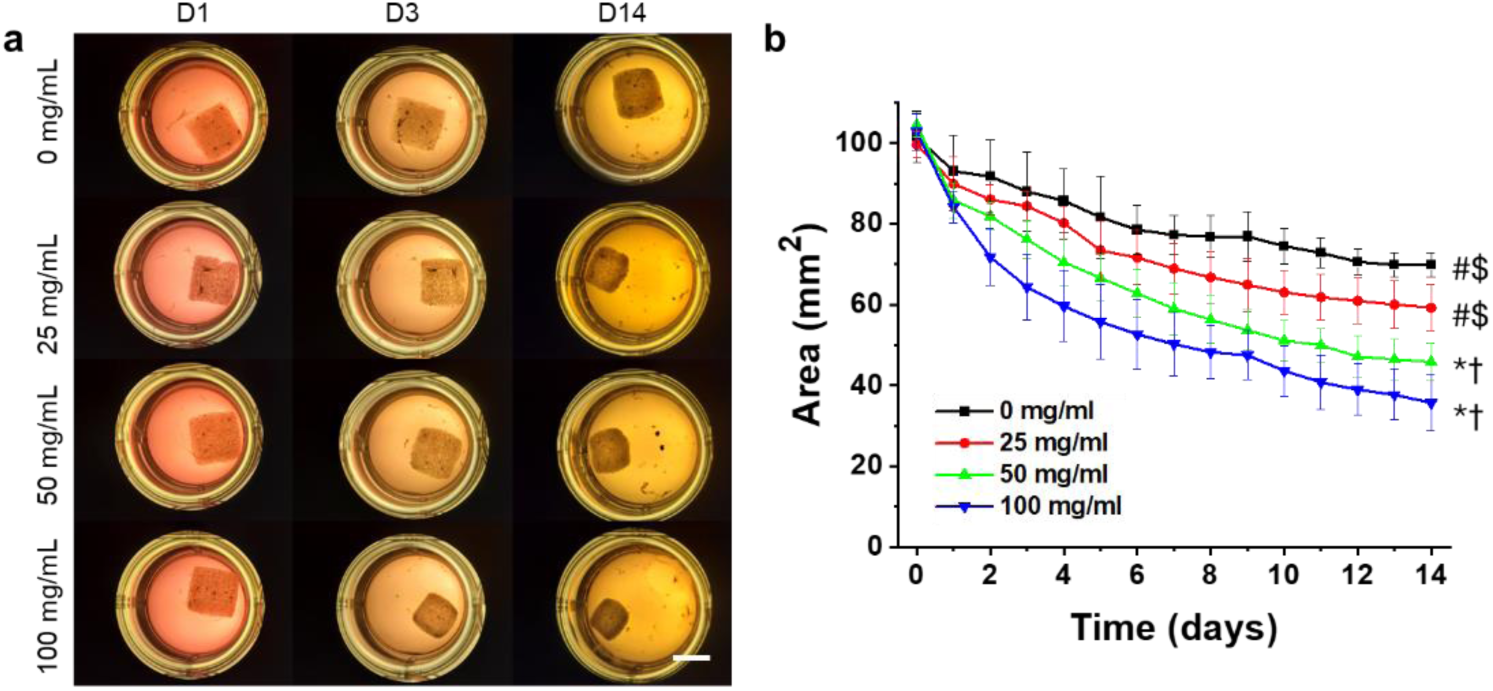
Shrinkage of 3D-printed cell-laden constructs. (a) OMA/GelMA composite bioinks were loaded with varying concentrations of gelatin microspheres. One mL of this bioink was added to 100 million cells to form the cell-laden bioink. Here, the bioink was used to print 10 mm × 10 mm × 0.6 mm squares, crosslinked directly after printing with UV light, and then monitored for 14 days to evaluate shrinkage. A selection of images showed the progression from initial to final shape over time. (b) Quantifying the area of the constructs over time revealed that increasing gelatin microsphere concentration led to an increase in shrinkage. Scale bar = 10 mm. “*”, “†”, “#”, and “$” indicate statistical significance at day 14 compared to 0, 25, 50, and 100 mg/mL, respectively (p < 0.05).

To investigate the ability to control the direction of bending in our 4D system, a bilayer approach was then pursued. Here, the bottom layer consists only of OMA microgels while the top layer consists of the cell-laden composite hydrogel. The top layer was printed directly onto the bottom layer and the entire construct was photocrosslinked to obtain the desired bilayer structure. Cells within the top layer made physical connections with each other and the surrounding biomaterial matrix via cell adhesions. This enabled cytoskeleton-generated CCF in the cell-laden layer to cause macroscopic shrinkage of this layer. The OMA microgel-only layer resisted this contraction, causing the bilayer to bend towards the cell-laden layer direction, as shown in the cell growth medium (GM) condition (**Fig. 3a, GM**). In contrast, cell-free bilayers cultured under the same conditions demonstrated negligible shape changes over a 14-day period (**Fig. S4**). To validate the role of CCF in driving the observed construct geometric changes, bilayer constructs were cultured in GM containing Cytochalasin D (CytoD), a known inhibitor of actin polymerization. Since cellular forces are generated by the activation of the actin-myosin complex^38^, inhibiting actin polymerization greatly diminishes a cell’s ability to generate forces. Here, CytoD was supplemented in growth medium at a concentration of 5 µM, which has previously been shown in the literature to disrupt the cytoskeleton by preventing F-actin formation and inhibiting cell motility ^39, 40, 41^. Ultimately, it was expected that CytoD would prevent the cells from producing CCF, thereby in turn inhibiting shape transformations. Constructs cultured in this media displayed no macroscopic changes in shape over the duration of culture (**Fig. 3b, Cyto D**) and no sign of cell death at day 14 (**Fig. S5**). Since CytoD was dissolved in dimethyl sulfoxide (DMSO) and then added to GM, an additional group of GM containing 0.1% v/v DMSO was included. To corroborate that this concentration of DMSO had no effect on bending, a vehicle control condition was established in which constructs were cultured in growth medium with 0.1% DMSO (**Fig. 3c**). The normal GM and DMSO conditions showed no significant differences in bending angle over 14 days. The similar bending angles quantified from constructs culured GM and DMSO indicated comparable hydrogel deformations (**Fig. 3b**). Histological analysis was performed to investigate the microscale effects of CytoD treatment (**Fig. 3c**). Hematoxylin and eosin (H&E) staining of samples in each media condition illustrated that cells in the GM and DMSO groups appeared to have elongated cell bodies and were able to form a fibrous border along the outer surfaces of the cell-laden layers, consistent with normal behavior of fibroblasts^42^. In contrast, cells in the CytoD group displayed predominantly rounded cell bodies and were unable to form a fibrous border, consistent with the inhibition mechanism of this drug^39, 40^. These results suggest that deformation was ascribed to the cell-mediated matrix contraction of the upper (cell-laden) layer. To the best of our knowledge, this is the first time that controllable matrix shape transformation replying soly on CCF was demonstrated in a uniformly loaded cell construct printed using a composite bioink with high free-standing stability.

**Fig. 3.**
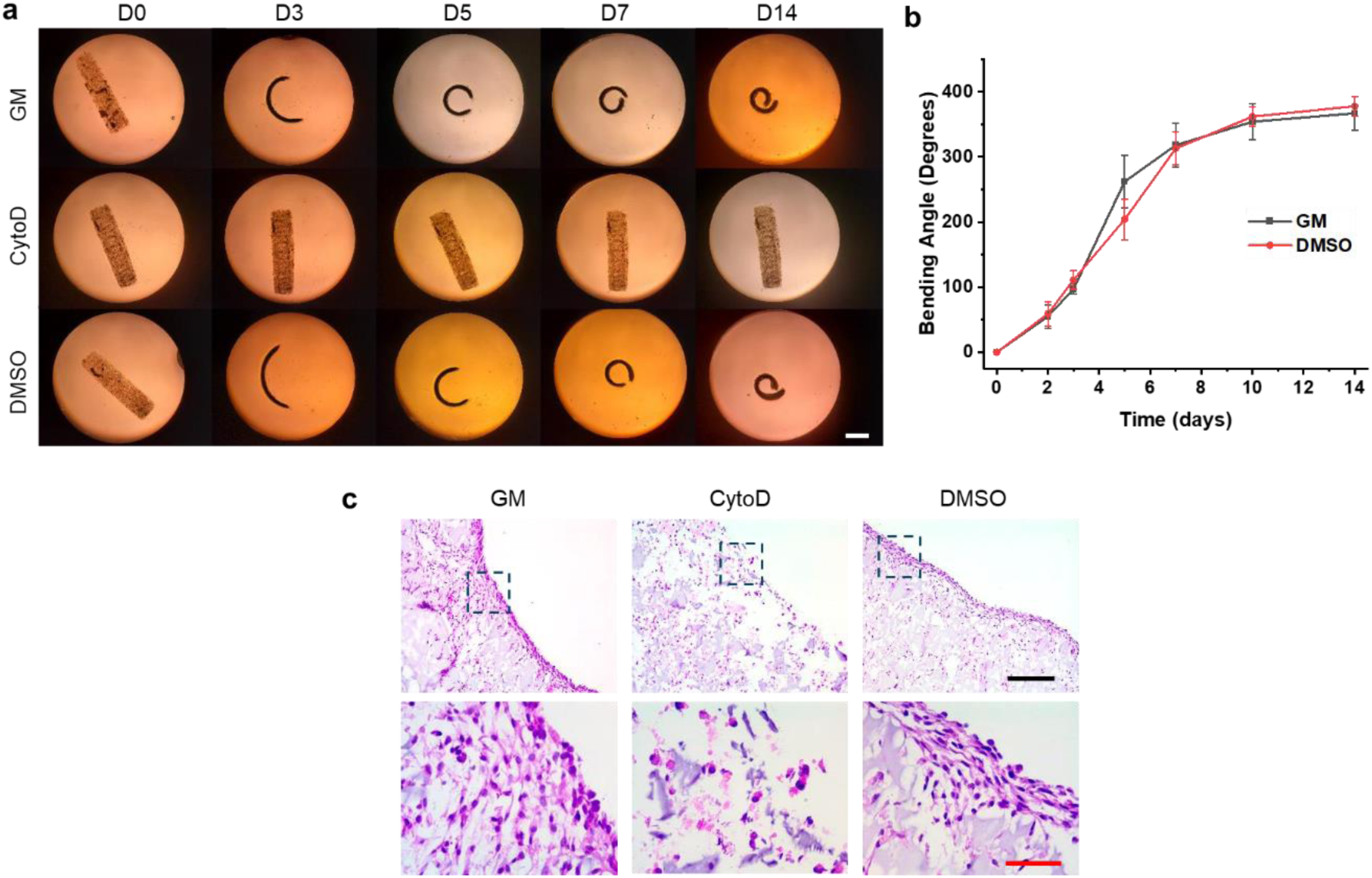
4D shape morphing in bilayer constructs. (a) To examine the role of CCF in the 4D process, printed constructs were cultured in cell growth media (GM) for 14 days. Constructs were also cultured in GM containing 5 µM CytoD as a negative control, and in GM containing 0.1% DMSO as a vehicle control. (b) Constructs in normal GM and GM with 0.1% DMSO conditions exhibited similar 4D bending, demonstrating that CytoD was responsible for the lack of bending seen in this condition. The average bending angles in the GM and DMSO conditions were not significantly different from each other at all timepoints (p > 0.05). (c) Photomicrographs of histological staining with H&E at day 14 showed cells in GM and DMSO conditions forming fibrous borders while cells in Cyto D condition did not. White scale bar = 5 mm. Black scale bar = 200 µm. Red scale bar = 50 µm.

Previous studies have shown that the scaffolds seeded with chondrocytes exhibit more pronounced contraction compared to those seeded with MSCs^43, 44^. Therefore, it is expected that the hydrogels containing hMSCs could undergo enhanced deformation when exposed to chondrogenic differentitaion conditions. Furthermore, coordinating cell differentiation and new cartilage tissue formation with cell-mediated 4D construct shape transformations could advance biomimetic tissue engineering. To explore this, we investigated the effects of chondrogenic differentiation of hMSCs within the cell-laden layers and extent of 4D geometric changes in the bilayered constructs. To accomplish this, hMSCs at a density of 1 × 10^8^ cells/mL bioink were suspended in the composite bioink and bilayer constructs were printed as designed in **Scheme 1d**. Constructs were cultured in one of two media: (1) normal GM, or (2) chondrogenic pellet medium (CPM), a serum-free medium which includes TGF-β1. Construct bending angles were measured from photomicrographs of constructs in each condition over 21 days of culture (**Fig. 4a**). The bending angles of constructs increased progressively during culture, with constructs in CPM exhibiting significantly greater (p < 0.05) bending than those in GM on days 3, 5, 7 and 14 (**Fig. 4b**). By day 21, constructs in both GM and CPM displayed comparable bending angles, indicating that exposure to chondrogenic factors induced a greater rate of bending in hMSC-laden constructs but did not increase the maximum bending angle achieved by day 21.

**Fig. 4.**
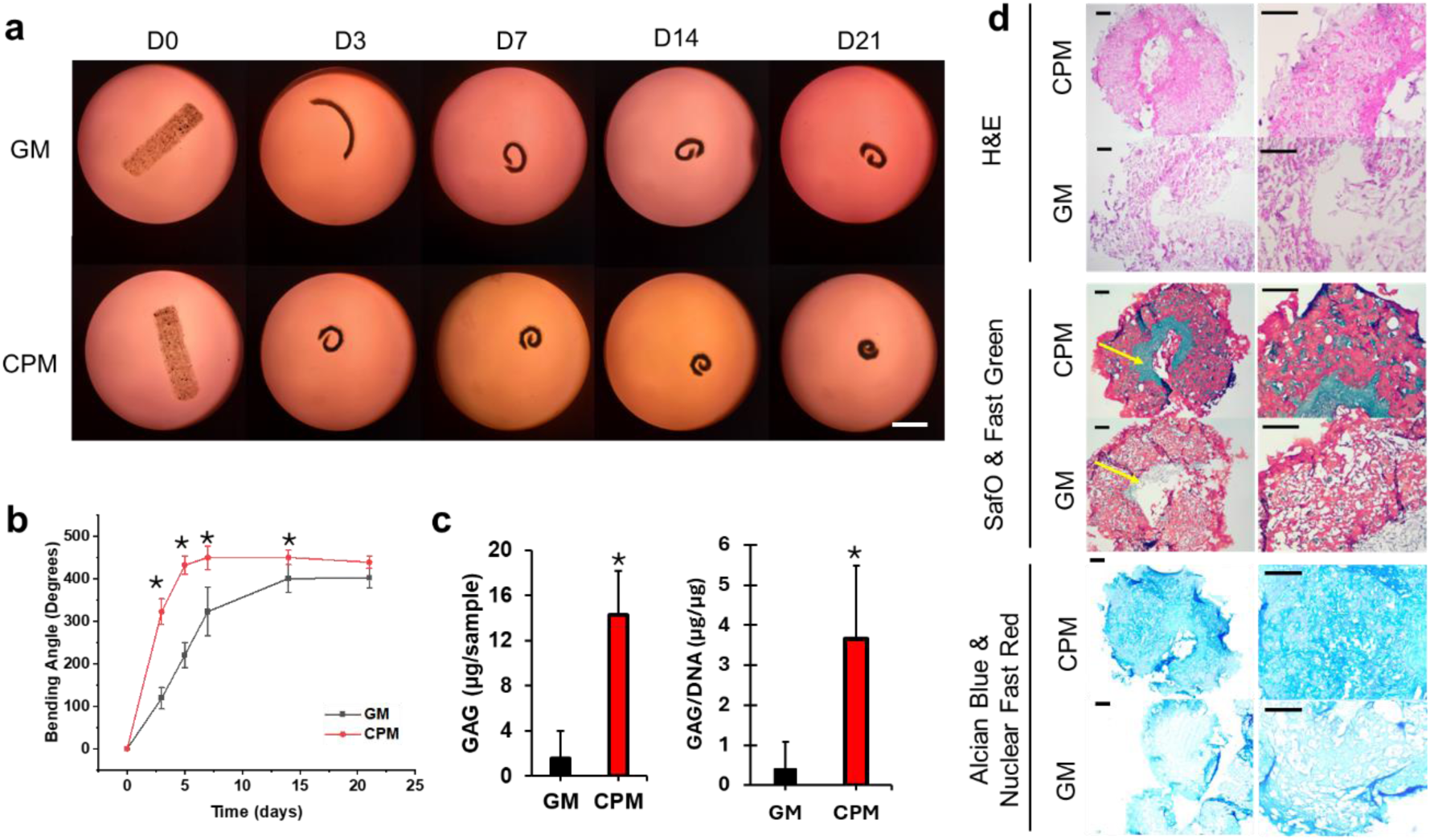
Chondrogenesis in 4D bilayer constructs. hMSCs were printed in bilayer constructs and cultured in normal growth media (GM) or chondrogenic pellet medium (CPM) and for 21 days. (a) Macroscopic images of constructs cultured over 21 days. (b) Quantification of bending angle over time (*N* = 4). (c) Quantification of GAG production and GAG/DNA levels for both GM and CPM conditions. Constructs cultured in CPM produced significantly more GAG (p < 0.05). (d) Histological staining on day 21 qualitatively confirmed substantial GAG production in constructs cultured in CPM.

During chondrogenic differentiation, hMSCs secrete a matrix rich in negatively charged polysaccharides known as glycosaminoglycans (GAGs), which contribute to the unique mechanical properties of cartilage^45^. Therefore, the presence of GAGs is a useful measure to evaulate the extent of chondrogenesis in the bilayer constructs cultured in GM and CPM media conditions. Biochemical analysis revealed that constructs cultured in CPM had significantly higher (p < 0.05) GAG content normalized to DNA content, consistent with an increase in chondrogenesis (**Fig. 4c**). Additionally, histological analysis was performed to visually corroborate the biochemical results (**Fig. 4d**). H&E staining revealed that constructs cultured in CPM formed a more dense and compact tissue early in culture compared to the those cultured in GM. This suggests that chondrogenic differentiation of hMSCs resulted in a higher degree of condensation and, consequently, generated a higher local cell density and enhanced cell-cell interactions than those in GM, leading to a more pronounced bending observed visually. Staining for Safranin O and Fast Green revealed the presence of negatively charged GAGs and collagen, respectively. In both CPM- and GM-cultured constructs, much of the collagen staining was localized to the surface of cell-laden layer directly in contact with media (inner side, as indicated by the yellow arrows), consistent with previous results where cells from cell aggregates in direct contact to the media often produce a fibrous layer (outer side) during chondrogenic differentiation^46, 47, 48^. While the OMA matrix in the construct can also be nonspecifically stained by Safranin O, CPM-cultured constructs exhibited more intense Safranin O staining within the inner bounds of the cell-laden layer, which supports the increase in normalized GAG levels measured in the biochemical analysis. To further substantiate this observation, constructs were additionally stained with Alcian blue at a pH of 0.2, which selectively stains strongly sulphated proteoglycans from cell-produced GAG without staining the OMA microgels or GelMA. The lack of intense blue staining in the GM-cultured constructs showed that the majority of Safranin O staining is due to the presence of OMA and not cell-produced GAG. Additionally, the intense blue staining of the CPM-cultured constructs revealed that much of the Safranin O staining is indeed cell-produced GAG. These results demonstrate that cells subject to chondrogenic differentiation culture conditions exhibited amplified and accelerated initial construct morphing. Additionally, the encapsulated cells were able to produce a cartilage-like matrix. Therefore, this system exhibits immense potential for CCF-based 4D chondrogenic tissue regeneration.

Next, it was investigated whether the direction of contraction can be controlled by patterning the cell-laden layer to induce complex 4D events (**Fig. 5a**). For example, printing parallel lines of cell-laden hydrogel on a rectangular hydrogel layer resulted in the formation of a cylindrical tube-like structure. Similarly, printing parallel cell-laden lines diagonally across a rectangular hydrogel layer resulted in the formation of a helical structure. Using 3D printing to create these structures also allowed for the generation of 4D constructs that change shape along multiple axes simultaneously. To illustrate this, a 4-armed bilayer “gripper” shape was printed. Each arm was observed to bend upward and inward, similar to how one’s fingers bend to grip an object in one’s palm. Additionally, a bilayer rectangle was printed with the cell-laden layer on top and adjoined to a second bilayer rectangle printed cell-laden layer first. Accordingly, bending was observed in opposite centrosymmetric directions around same axis, resulting in an “S”-shaped structure. These results indicate that cellular forces were sufficient to drive spatiotemporal changes within this biopolymer microenvironment, and that these changes persisted over 14 days of culture. Importantly, the complex final structures were generated and controlled by the precise patterning of the printed hydrogel and cell-laden layers.

**Fig. 5.**
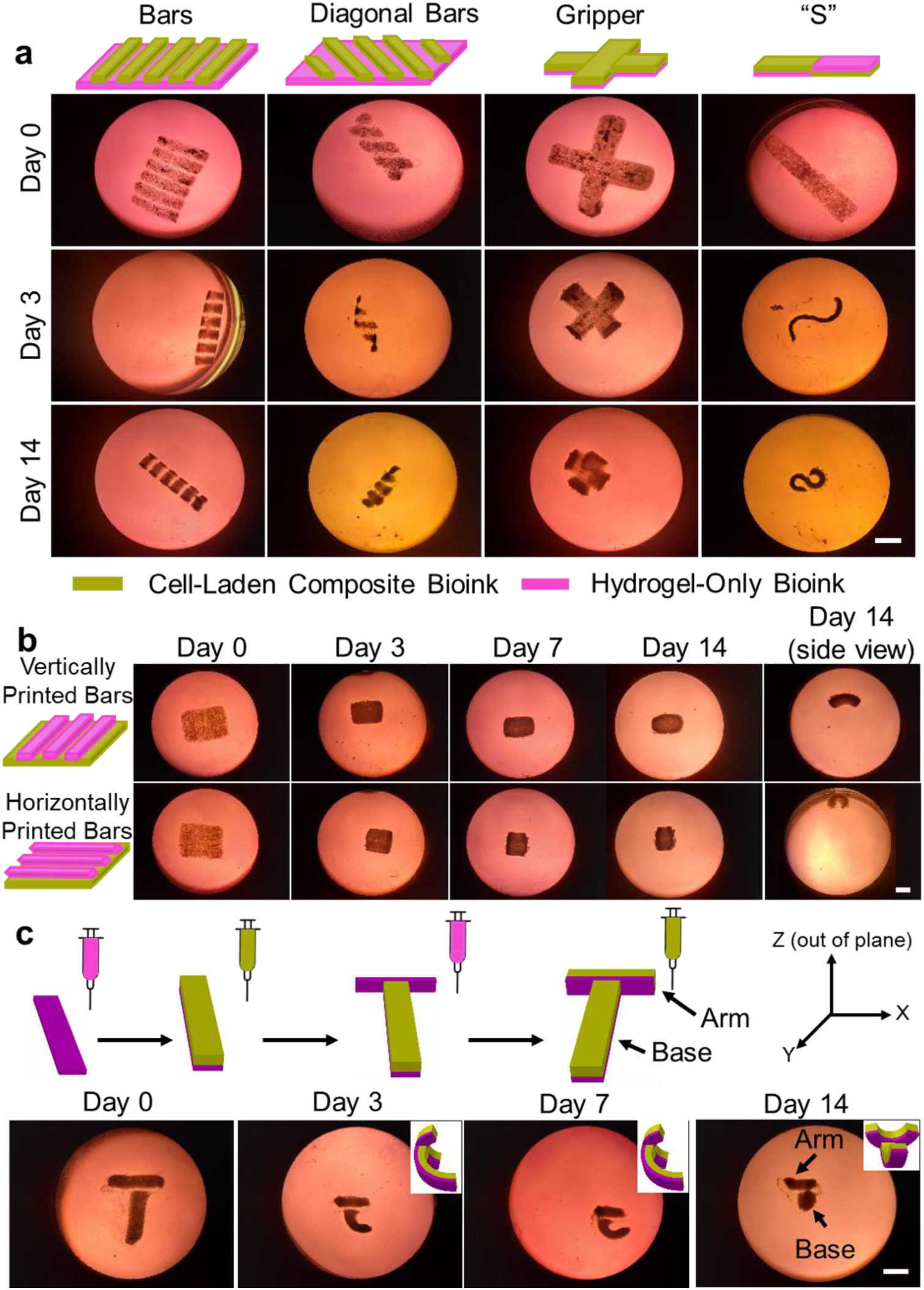
Complex patterning of the cell-laden and cell-free hydrogel layers. (a) Complex patterning of the cell-laden layer. Schematics of the printed constructs followed by photomicrographs at days 0, 3 and 14. (b) Complex patterning of the cell-free hydrogel layer. Rectangular hydrogel patterns were printed either vertically or horizontally onto a cell-laden hydrogel rectangle. The direction of the hydrogel pattern influenced the bending direction of the constructs. (c) Complex patterning of both the cell-laden and cell-free hydrogel layers. Layers were printed sequentially according to the schematic, resulting in bending around two separate, non-parallel axes. Scale bars = 10mm

Up to this point, directional bending has been controlled solely by patterning the cell-laden layer. Therefore, we investigated whether the direction of 4D bending could be influenced by patterning the cell-free layer. To demonstrate this, the composite cell-laden bioink was printed into 9 mm × 12 mm × 0.6 mm rectangles. A patterned cell-free hydrogel layer consisting of either horizontal or vertical bars was then printed onto the composite cell-laden layer. The cell-free hydrogel bars were observed to induce bending of the constructs around an axis perpendicular to the direction of the bars (**Fig. 5b**). Differences in initial printed geometries (e.g., rotated 90 degrees) contributed to differences in shape change over the duration of the 14-day culture period.

In addition to controlling the direction of contraction by separately patterning the cell-free and cell-laden layers, multiple bending axes can be incorporated into a single construct by patterning both layers simultaneously. To demonstrate this, T-shaped bilayer structures were printed in a step-by-step protocol. **Fig. 5c** presents the schematic of printing, where the base of the T was first formed by printing a rectangular cell-free hydrogel layer, followed by a matching cell-laden layer directly on top. Next, the arm of the T was formed by printing a cell-free hydrogel layer directly perpendicular to the base and subsequently printing a cell-laden layer adjacent to the perpendicular cell-free hydrogel layer. This initial geometry caused the arm of the T to bend around the y-axis, while the base of the T exhibited bending around the x-axis. Multi-axial bending around two non-parallel axes has not previously been demonstrated using generated CCF in cell-laden biomaterials.

## 3. Discussion

Controlling shape transformation in 4D reconstituted tissues has garnered significant attention in tissue regeneration. This capability is critical as it allows engineered tissues to recapitulate essential morphogenetic processes that occur *in vivo*, providing a powerful tool to mimic natural tissue development^49^. The programming and control of shape morphing in engineered living constructs are fundamental for advancing tissue engineering. Fortunately, emerging 4D tissue bioprinting^50^, which produces shape-transformable objects, offers a promising solution to push this frontier forward.

However, 4D tissue bioprinting faces critical challenges due to the demanding requirements for cytocompatible materials for cell embedding, long-term culture, and cytocompatible stimuli that permit morphing under physiological conditions. Consequently, recent developments have focused on identifying suitable biomaterials. Typically, engineered constructs based on these biomaterials undergo shape morphing in response to external stimuli such as light, magnetic fields, and solvents. CCF plays a crucial role in cell differentiation, migration, and growth during development, tissue remodeling and healing. It also plays a vital role in morphogenesis by driving tissue bending, stretching, alignment, and repositioning^51^. Thus, developing a printable system capable of promoting controllable construct shape transformation on a large scale in response to internal CCF is highly significant, as it enables the creation of structurally dynamic tissues. However, the utilization of inherent CCF generated by cells within artificial tissues has been scarcely reported.

Current systems for this purpose are based on morphing cell condensates and cell-laden scaffolds, which are limited to microscale size, local morphing, or ultra-soft biomaterials. With this in mind, in this work we developed a system with bioink mechanical properties robust enough for printing stand-alone constructs while allowing for transformation by weak CCF generated by uniformly distributed cells throughout the matrix. We achieved this by developing printable jammed OMA microgel bioinks and incorporating GelMA to enhance cell adhesion and interactions. Importantly, we devised sacrificial gelatin microspheres to transition the mechanically robust printed and photocrosslinked hydrogel scaffold to a mechanically soft state during culture while maintaining structural integrity. To control spatial shape change, an inert thin OMA hydrogel layer was used as a CCF resistant layer. As a result, the bilayer composed of an OMA layer and a cell-laden composite hydrogel layer exhibited predefined deformation. With this system, we generated complex shape transformations by carefully arranging the materials’ spatial positions with the aid of 3D printing without the need of a support structure or slurry bath. Additionally, when subjected to a differentiation environment, cartilaginous tissue was generated while promoting CCF-mediated tissue reshaping. Thus, this system, demonstrating CCF dynamism, marks a significant step towards mimicking *in vivo* tissue development, signifies a major advancement in 4D technology, and has the potential to profoundly impact 4D tissue engineering. Furthermore, by demonstrating the ability to generate complex curvatures, this system can effectively emulate some intricate tissue morphogenesis observed during native tissue development. Future research could leverage this system as an advanced tool for normal and diseased tissue modeling by incorporating cell-directed tissue dynamics in addition to the bulk properties typically controlled in traditional 3D models. Additionally, this platform has potential applications in the development of soft biohyrid robotics^52^ that utilize living cell actuation, offering advantages such as enhanced energy efficiency and autonomous motion. Another promising direction for future work includes the design of self-driven cargo or drug delivery systems that operate independently of external energy input.

However, the current system also faces challenges, including: (1) low construct resolution due to the inherent limitations of extrusion printing; (2) multimodal shape transformations demonstrated in a single construct are still not complex enough to mimic many intricate shape changes occurring *in vivo*. Future developments must address these limitations. The rapid progress of this emerging technology is expected to have a profound impact on regenerative medicine, biomedical engineering, tissue engineering, and beyond.

In summary, a novel bioink composed of OMA microgels, GelMA, gelatin microspheres, and living cells has been developed for 4D bioprinting of living constructs that undergo controllable spatiotemporal geometric changes mediated by cell-generated forces, i.e., CCF. By forming bilayered constructs, we demonstrated that the direction of 4D bending can be controlled. Multiple degrees and directions of deformation were observed within a single construct, showing the capacity for complex shape transformations. hMSCs can be encapsulated in the living constructs at high cell density, with the supplementation of chondrogenic signals leading to an increased rate of 4D bending while simultaneously guiding cells down the chondrogenic lineage. Additionally, each layer of the bilayered constructs can be spatially arranged with the aid of 3D printing to generate complex initial and final structures. Both the cell-free and cell-laden layers can be patterned to provide new shape transformations in culture, illustrating the system’s versatility. Our system provides the first report of CCF-mediated 4D bioprinted living constructs that demonstrate highly stable, complex shape transformation in a preprogrammed and controlled fashion.

## 4. Methods

### Cell culture

NIH 3T3 (ATCC) were cultured in low-glucose Dulbecco’s modified eagle medium (DMEM, Sigma, cat#D5523-50L) supplemented with 10% fetal bovine serum (FBS, Sigma, cat#18N103) and 1% penicillin/streptomycin (P/S, Gibco, cat#15140122). Upon reaching 90% confluence, cells were trypsinized, counted, and pelleted at 100 million cells per vial to be combined with 1 mL of composite bioink. After printing, constructs were cultured in high-glucose DMEM (Sigma, cat# D5648-10x1L) supplemented with 10% FBS and 1% P/S.

### OMA Microgel Synthesis

Alginate was modified with 2% oxidation and 30% methacrylation according to previously published protocols^53^. Briefly, 1% sodium alginate (10 g, Protanal LF 20/40, FMC Biopolymer) solution was dissolved in ultrapure deionized water (diH_2_O, 900 mL) by stirring overnight at room temperature (RT). 216 mg of sodium periodate was dissolved in 100 mL of diH_2_O, mixed with the alginate solution to achieve 2% theoretical alginate oxidation and reacted in the dark at RT for 24 hrs under stirring. 2-morpholinoethanesulfonic acid (MES, 19.52 g, Sigma) and sodium chloride (NaCl,17.53 g, Sigma) were then dissolved in the oxidized alginate solution and the pH was adjusted to 6.5 using 4 N sodium hydroxide (NaOH, Sigma). N-hydroxysuccinimide (NHS, 1.77 g, Sigma) and 1-ethyl-3-(3-dimethylaminopropyl)-carbodiimide hydrochloride (EDC, 5.84 g, Sigma) were dissolved into the mixture. AEMA (2.54 g, Polysciences) was then slowly added to the solution to achieve a theoretical methacrylation level of 30%. The reaction was conducted at RT for 24 hrs in the dark. The reacted OMA solution then was poured into excess acetone to precipitate the OMA. The precipitate was dried in a fume hood and subsequently dissolved in diH_2_O at a 1% w/v concentration. The OMA solution was dialyzed for purification using a dialysis membrane (MWCO 3500, Spectrum Laboratories Inc.) for 3 days. The dialyzed OMA solution was collected and treated with activated charcoal (5 g/L, 50-200 mesh, Fisher) for 30 min. The solution was further purified and sterilized by filtering through a 0.22 μm pore membrane and then lyophilized. The ^1^H NMR characterization of the OMA is shown in **Fig. S6**. The actual methacrylation degree was determined to be 9.9% through ^1^H NMR analysis, as described in literature^54^.

Dry OMA was dissolved in MilliQ water to form a 2% w/v solution. The solution was then added dropwise to a beaker of 0.2 M calcium chloride under vigorous stirring and allowed to ionically crosslink for 4 hrs. Crosslinked OMA was collected and placed in a blender (Oster) with 100 mL of 70% ethanol. OMA was blended for two min before adding 50 mL of 70% ethanol, and then blending was continued for 2 min. OMA microgels and ethanol were collected into 50 mL conical tubes, centrifuged at 4200 rpm (Sorvall ST40R, thermofisher) for 5 min at 20 °C, and stored at 4 °C for future use.

### GelMA Synthesis

GelMA was synthesized according to previously established protocols^55, 56^. Briefly, 10 g of gelatin (type A, Sigma Aldrich) was dissolved in 100 mL of PBS (pH 7.4) and heated to 50 °C. Then 10 mL of methacrylic anhydride was added into the 10% gelatin solution and reacted for 1 hr at 50 °C and then stirred overnight at RT. GelMA was precipitated with acetone, purified via dialysis at 50 °C for 7 days with a MWCO 12-14k membrane (Spectrum Laboratories Inc.), sterilized via a 0.22 mm pore filter, and then lyophilized. The ^1^H NMR characterization of the GelMA is shown in **Fig. S7**. The actual methacrylation degree was determined to be 84% through ^1^H NMR analysis, as described in literature^54^.

### Gelatin Microsphere Synthesis

To synthesize gelatin microspheres, 10 g of gelatin type A (Sigma-Aldrich, cat#G2000-500G) was dissolved in 90 mL of diH_2_O (11% w/v) by heating at 60 °C. The gelatin solution was then added dropwise into 500 mL of preheated olive oil (45 °C) under vigorous stirring at a rate of 10 mL/min. After 10 min, the temperature was reduced to 15 °C while maintaining stirring. After 30 min, 200 mL of chilled acetone was added to the emulsion, and the mixture was stirred for 5 min. The gelatin microparticles were then collected by filtration and washed five times with acetone to remove any residual olive oil. Finally, the microspheres were dried in a fume hood overnight. The as-synthesized gelatin microspheres had an average size of 67 ± 34 μm, as shown in **Fig. S8**.

### Cell-Free Bioink and Cell-Laden Composite Bioink Preparation

The cell-free bioink is composed solely of OMA microgels, which were prepared as follows: OMA microgels were reconstituted through three washes of 0.05% photoinitiator (2-hydroxy-4’-(2-hydroxyethoxy)-2-methylpropiophenone, Sigma, PI)-containing MilliQ water and two washes of low-glucose DMEM with 0.05% PI. To prepare the cell-laden composite bioink, lyophilized GelMA was weighed and dissolved directly in the above reconstituted OMA microgels (3% w/v GelMA). Lyophilized gelatin microspheres were weighed and rehydrated for 15 min in low-glucose DMEM with 0.05% PI at a ratio of 15 µL/mg. The desired volume of OMA/GelMA mixture was measured and added to the rehydrated gelatin microspheres. One mL of this solution was added to a pellet of 100 million cells to form the cell-laden bioink.

### Single Layer Printing

A 3 mL syringe with a 22 gauge needle was loaded with cell-laden composite bioink and placed in a Cellink BIOX3D printer (Celllink, San Diego, CA). Square constructs measuring 10 mm × 10 mm × 0.6 mm were printed from a custom-made STL file using a print speed of 4 mm/s, an extrusion rate of 1.2 µL/s, and an infill density of 60%. Printed constructs were crosslinked with UV light at an intensity of 12 mW/cm^2^ for 30 s and subsequently transferred to 6 well plates containing 8 mL of media. To minimize disturbance to the constructs during culture, half the media was removed and replenished every day. Constructs were imaged daily using a dissection microscope and a Samsung Galaxy S9 phone camera.

### Bilayer Printing

To accomplish bilayer printing, two syringes were loaded into the 3D printer: the first contained only reconstituted OMA microgels (cell-free bioink); the second contained the cell-laden composite bioink. For bilayer rectangles, a 25 mm × 25 mm × 0.2 mm square was printed with the cell-free bioink using a 4 mm/s print speed, 1.2 µL/s extrusion rate, and 80% infill density. Three cell-laden rectangles measuring 18 mm × 4 mm × 0.6 mm were then printed using the cell-laden composite bioinks directly on top of the cell-free square at parameters of 4 mm/s, an extrusion rate of 1.2 µL/s, and an infill density of 60%, spaced 4 mm apart. After crosslinking constructs with UV light at an intensity of 12 mW/cm^2^ for 30 s, the cell-free layer was cut with a razor blade to match the geometry of the cell-laden rectangles, forming bilayer rectangles (*N* = 4).

To print constructs with complex cell-laden geometires, similar protocols were followed. A large cell-free layer was printed, followed by a patterned cell-laden layer. If necessary, the cell-free layer geometry was cut to match the cell-laden layer. Constructs with dual layers were constructed by printing a rectangle with dimensions of 12 mm × 9 mm × 0.6 mm. The second layer was then printed directly on top of the first layer using the cell-laden composite bioink. With the cell-free bioinkprinted first with using the cell-free bioink-ladenbioink

### CytoD Treatment

To further illustrate the role of cell-generated forces in the observed shape changes, the effect of CytoD (Invitrogen, cat#PHZ1063), a known inhibitor of actin polymerization, on bilayered constructs composed of a cell-free and a cell-laden layer was studied. CytoD was weighed and dissolved in sterile DMSO at a concentration of 5 mM and stored at 4 °C. Bilayer rectangles were printed and cultured in GM supplemented with 0.1% v/v CytoD, resulting in a final CytoD concentration of 5 µM. Half of the media was replaced every day, with fresh CytoD added at 0.1% v/v each time. To ensure that any change in construct behavior resulted from CytoD treatment and not DMSO, four bilayer rectangles were cultured in GM with 0.1% v/v DMSO. These media conditions were also compared to bilayer rectangles cultured in normal culture media. A total of four samples were prepared for each test unless otherwise specified (*N* = 4).

### Chondrogenesis in 4D Printed Constructs

Human mesenchymal stem cells (hMSC) were harvested according to previously established protocols^57^. For this study, hMSCs passage 3 were cultured in low-glucose DMEM supplemented with 10% FBS, 1% P/S, and 10 ng/mL fibroblast growth factor-2 (R&D, cat# 233-FB-MTO). Upon reaching 80% confluence, cells were trypsinized, counted, and pelleted at 100 million cells per vial to be combined with 1 mL of composite bioink. After printing with passage 4 cells, constructs were cultured in high-glucose DMEM (Sigma, cat#D5648-10X1L) supplemented with 10% v/v ITS+ (Fisher, cat#CB40352), 1% v/v non essential amino acids (NEAA, Gibco cat#11140050), 1% v/v P/S, 100 mM sodium pyruvate (Fisher, cat#SH3023901),10^-7^ M dexamethosone (Sigma, cat#D4902-100mg), L-ascorbic acid phosphate (Wako USA, cat#013-12061), and 10 ng/mL TGF-β1 (Peprotech, cat#100-21-10UG). Samples were collected for biochemical analysis (*N* = 3) and histological analysis (*N* = 2) at 3 weeks.

### Histology

After culture duration was complete, samples were fixed overnight in 10% neutral buffered formalin at room temperature (*N* = 2). Samples were then dehydrated through one-hour incubations in 70% ethanol, 95% ethanol, 100% ethanol (Fisher, cat#1SF1C163), 1:1 volume ratio of ethanol and xylene, and two separate 100% xylenes (Fisher, cat#x3s-4). Samples were then submerged in molten paraffin (Epredia, cat#8336) for at least 24 hrs before being embedded in paraffin blocks. 5-µm sections were obtained from blocks using a Leica RM2255 microtome (Leica Biosystems, Nussloch, Germany) and mounted on microscope slides. Mayer S Hematoxylin (Fisher, cat#TA125MH) and Eosin-phloxine solution, alcoholic (Electron Microscopy services, cat#26763-03) (H&E) sections were stained with hematoxylin for 2 min followed by counterstaining with eosin for 30 s. SafO (Acros Organics, cat#477-73-6) and Fast Green (Fisher, cat#F99-10) sections were stained with 0.1% w/v SafO for 5 min followed by counterstaining with 0.05% w/v Fast Green for 1 min. Alcian blue (Fisher, cat#92-31-9) and nuclear fast red (Polysciences, cat# 09773) sections were stained with alcian blue (pH = 0.2) for 30 min followed by counterstaining with nuclear fast red for 5 min.

### Biochemical Assays

DNA and GAG values were quantified according to previously described methods^57^. Briefly, GAG values were quantified by measuring the absorbance of DMMB-bound samples at 595 nm using a plate reader (*N* = 3). DNA values were similarly quantified by measuring fluorescence intensity of PicoGreen-bound samples at an excitation of 480 nm and emission of 520 nm (*N* = 3).

### Bending Angle Quantification

Dissection microscope images were used to measure bending angle as described in literature^18^. As shown in **Fig. S9**, a circle with crosshairs was superimposed on each image using Microsoft Powerpoint. The dimensions of the circle were modified to fit the arc of the construct in the image. The image and circle were then copied into ImageJ and the angle tool was used to measure the angle between one end of the construct, the intersection of the crosshairs, and the other end of the construct. Using this method, a construct that has bent into a half circle was measured as 180 degrees, while a construct whose ends are touching were measured as 360 degrees.

### Live/Dead Staining

Printed constructs were stained with fluorescein diacetate (Sigma, cat# F7378) and propidium iodide (Sigma, cat# P4170-25MG) to visualize live and dead cells, respectively. These stains were incubated with printed constructs for 5 min, after which all media was removed, and samples were immediately imaged using a Nikon Eclipse TE300 fluorescence microscope (Nikon, Tokyo, Japan) equipped with a AmScope MU1403 camera (AmScope, Irvine, California). *N* = 3.

### Statistics

All graphs are reported as mean ± standard deviation (±SD). For data with more than two groups, significance was determined using one-way ANOVA with a post-hoc Tukey HSD test. For data with only two groups, significance was determined using the Student’s t-test. p = 0.05 was considered significant unless otherwise specified.

## Acknowledgements

The authors gratefully acknowledge funding support from the Department of Veterans Affairs, Veterans Health Administration, Office of Research and Development, Rehabilitation Research and Development Service under award numbers RX004288 and RX004825 and the National Institutes of Health’s National Institute of Arthritis and Musculoskeletal and Skin Diseases under award number R01AR081448. The contents of this publication are solely the responsibility of the authors and do not necessarily represent the official views of the Department of Veterans Affairs or the National Institutes of Health.

## Author Contributions

**D.S.C**: Methodology, Investigation, Formal analysis, Visualization, Writing - Original Draft; **K.L.G**: Methodology, Investigation, Formal analysis, Visualization, Writing - Original Draft, Writing - Review & Editing. **A.D.**: Conceptualization, Methodology, Investigation, Formal analysis, Visualization, Writing - Original Draft, Writing - Review & Editing. **E.A.**: Conceptualization, Methodology, Formal analysis, Resources, Supervision, Funding acquisition, Writing - Review & Editing.

