## Supplemental Information for "Cell Contractile Forces Drive Spatiotemporal Morphing in 4D Bioprinted Living Constructs"

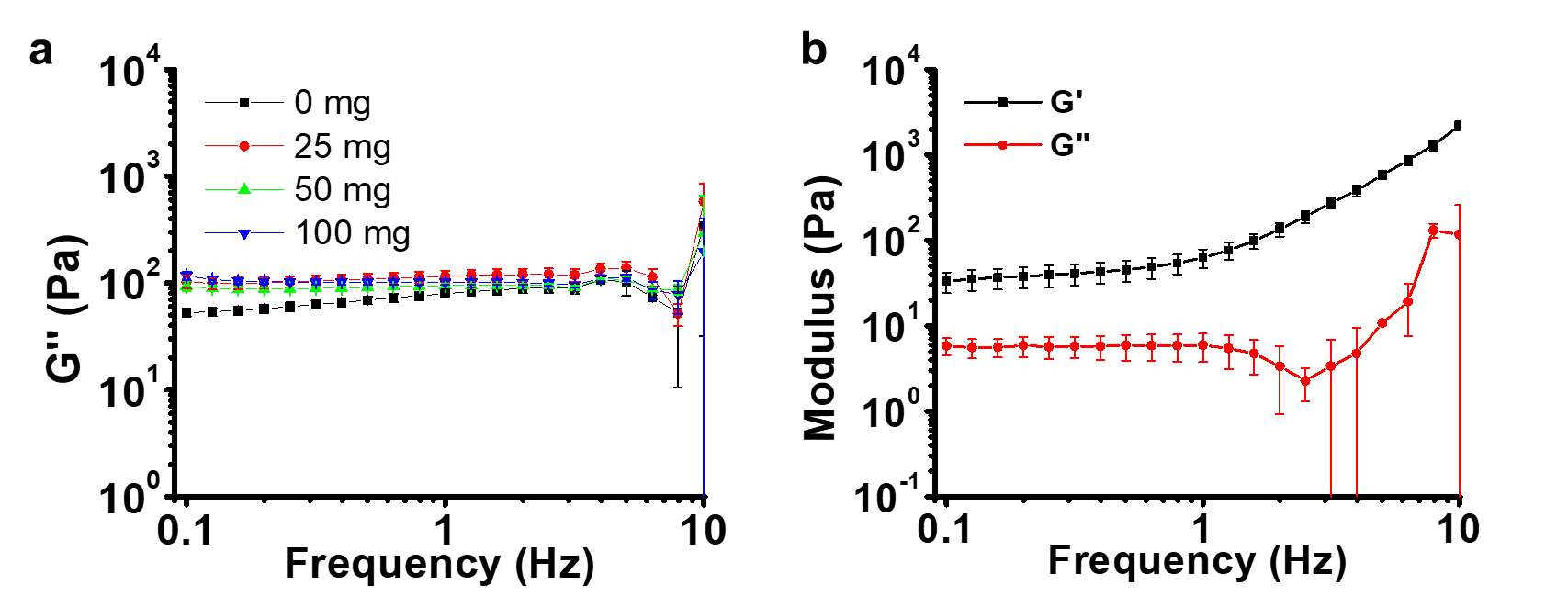

**Fig. S1**. (a) Loss modulus (G’’) of composite bioinks with varying gelatin microsphere concentrations of 0, 25, 50, and 100 mg/mL. (b) After one day of culture, the moduli of the composite bioinks containing 50 mg/mL gelatin microspheres have decreased but still maintain higher G’ than G’’.

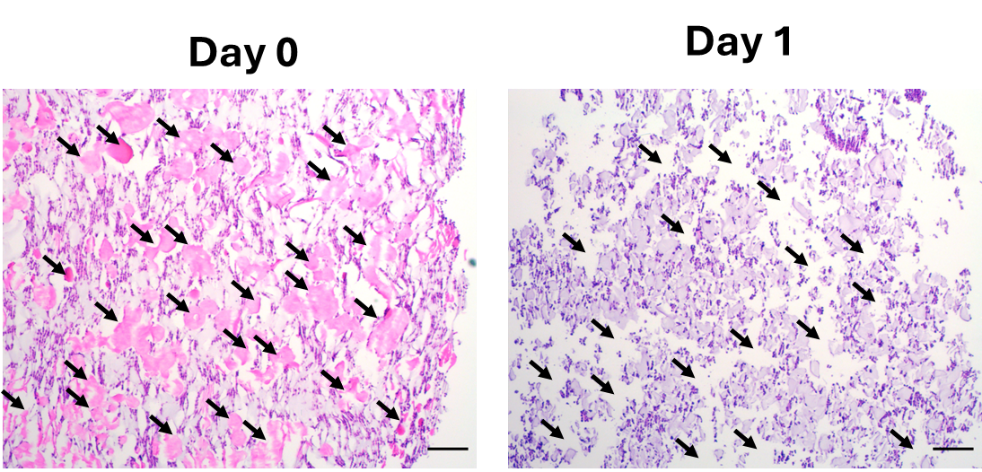

**Fig. S2**. Photomicrographs presenting histological evidence of the liquefactions of gelatin microspheres. On day 0, gelatin microspheres appeared light pink under H&E staining. After one day of culture, the gelatin had liquefied and diffused out of the constructs, leaving microscale pores (black arrows). Scale bars = 200 µm.

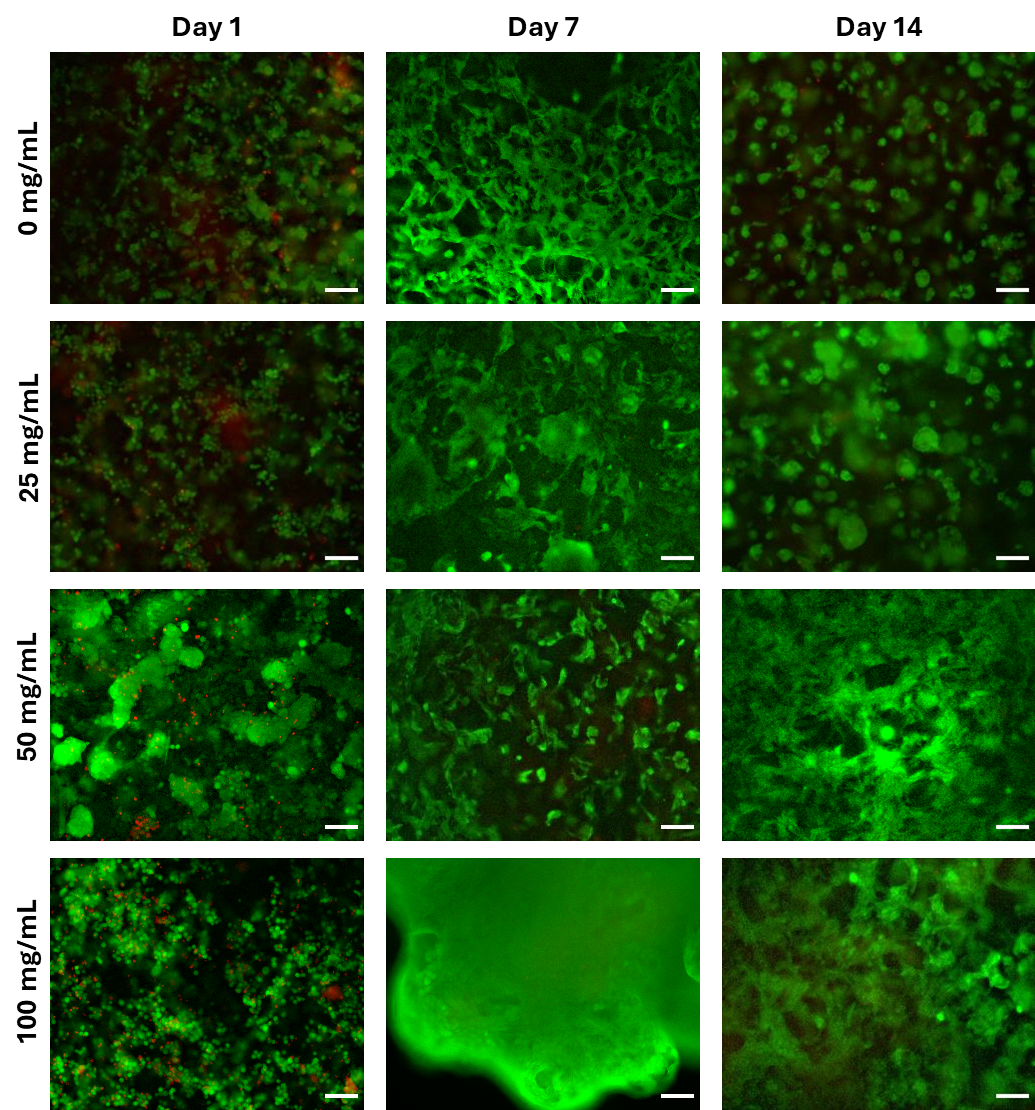

**Fig. S3**. Photomicrographs of live/dead staining showing high cell viability in printed constructs with varying concentrations of gelatin microspheres at days 0, 7 and 14. Scale bar = 100 µm.

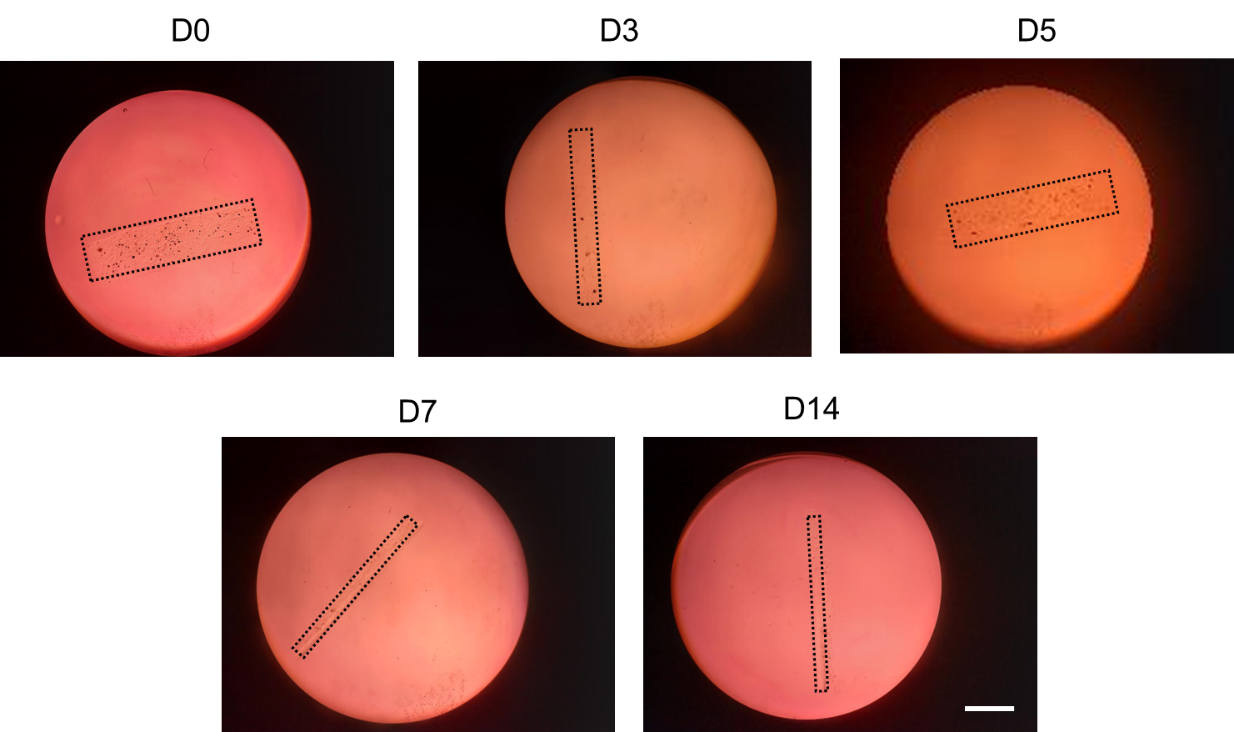

**Fig. S4**. Representative images showing the shape changes of cell-free bilayer strips over 14 days of culture in GM. Note that the visually different sizes of the bilayer strips in the images are due to variations in viewing angles. In the images of D0 and D5, the strips were imaged laying down in a “flattened” state; in the images of D7 and D14, the strips were imaged on their sides in “upright” state; and in the image of D3, the strip was imaged in an “inclined upright” state. Scale bar = 10 mm.

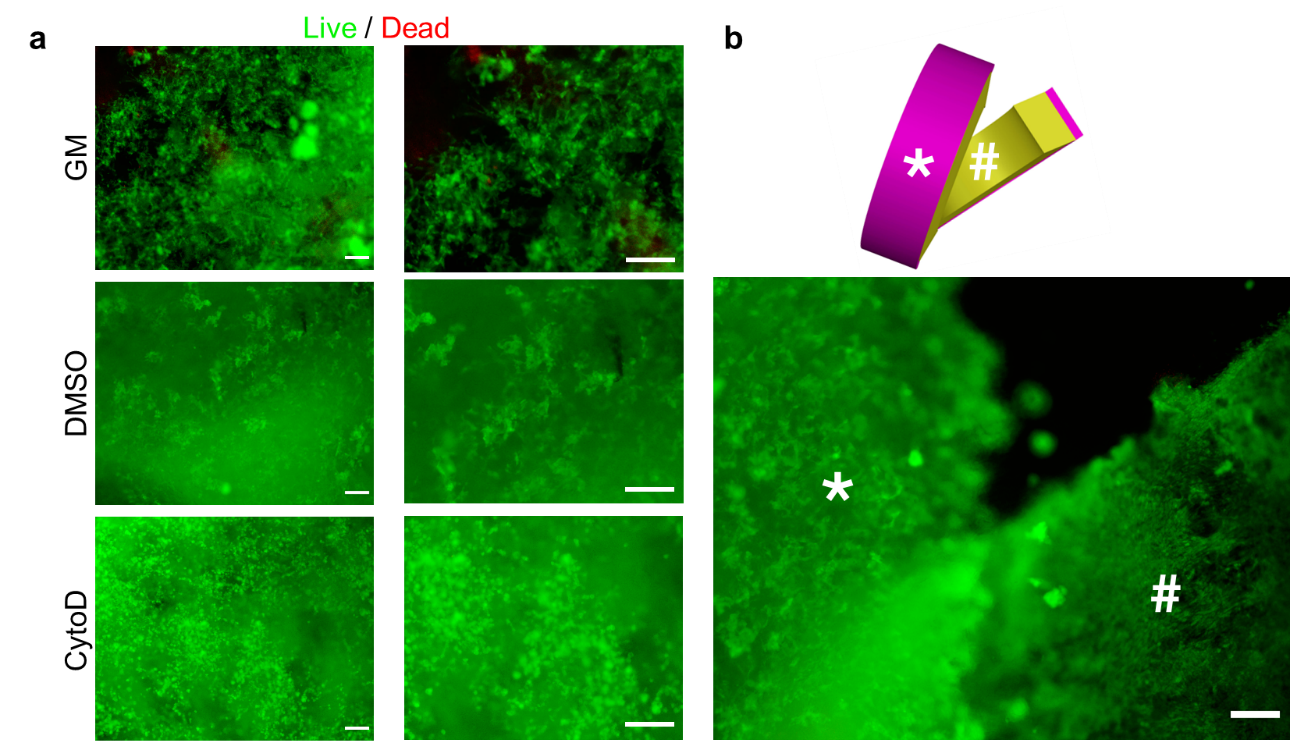

**Fig. S5**. Live/dead staining of cells constructs cultured in GM, DMSO and CytoD conditions on day 14. (a) Composite live/dead photomicrographs revealed that cell morphology appeared similar in GM and DMSO conditions, while cells appeared more rounded in the CytoD condition. (b) Photomicrograph of live/dead staining of a GM group construct, centered at the point where the construct layer in focus changes from the cell-free layer to the cell-laden layer. Imaging through the cell-free layer (indicated with *) shows high cell viability at the interface between the layers that is comparable to the cell-laden surface in contract with media (indicated with #). . Scale bars = 200 µm.

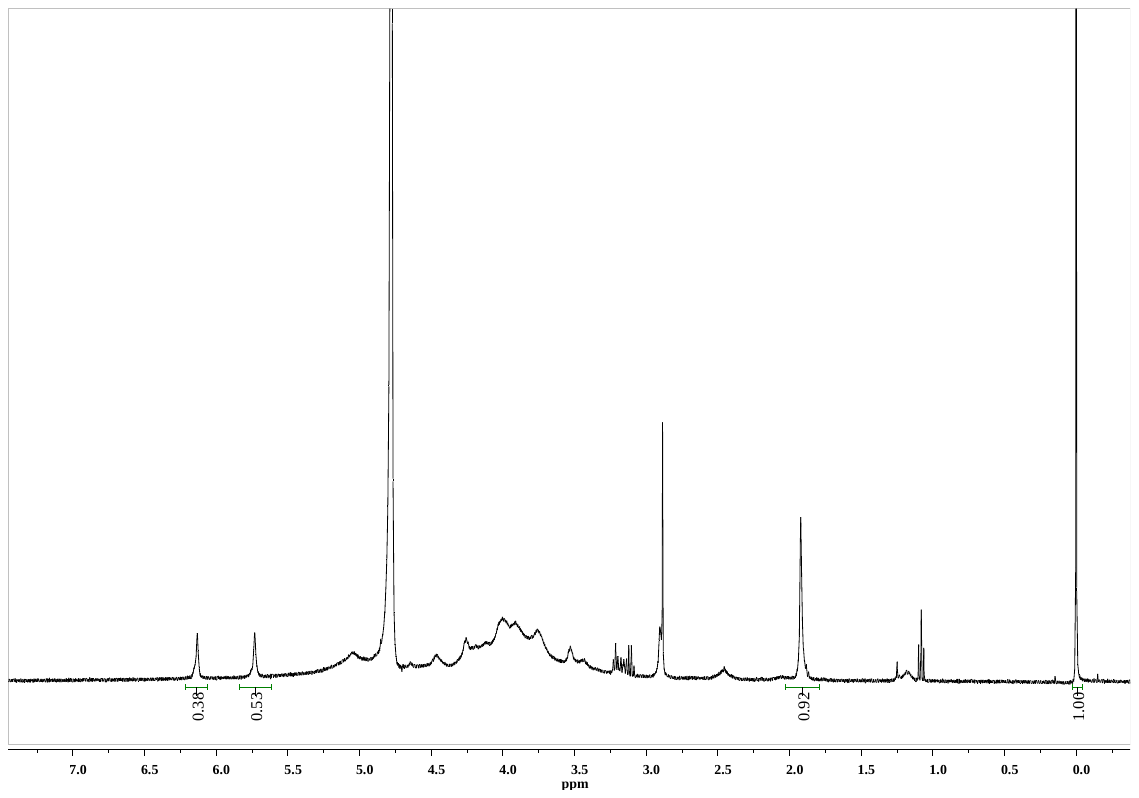

**Fig. S6**. ^1^H NMR spectrum of OMA in D_2_O.

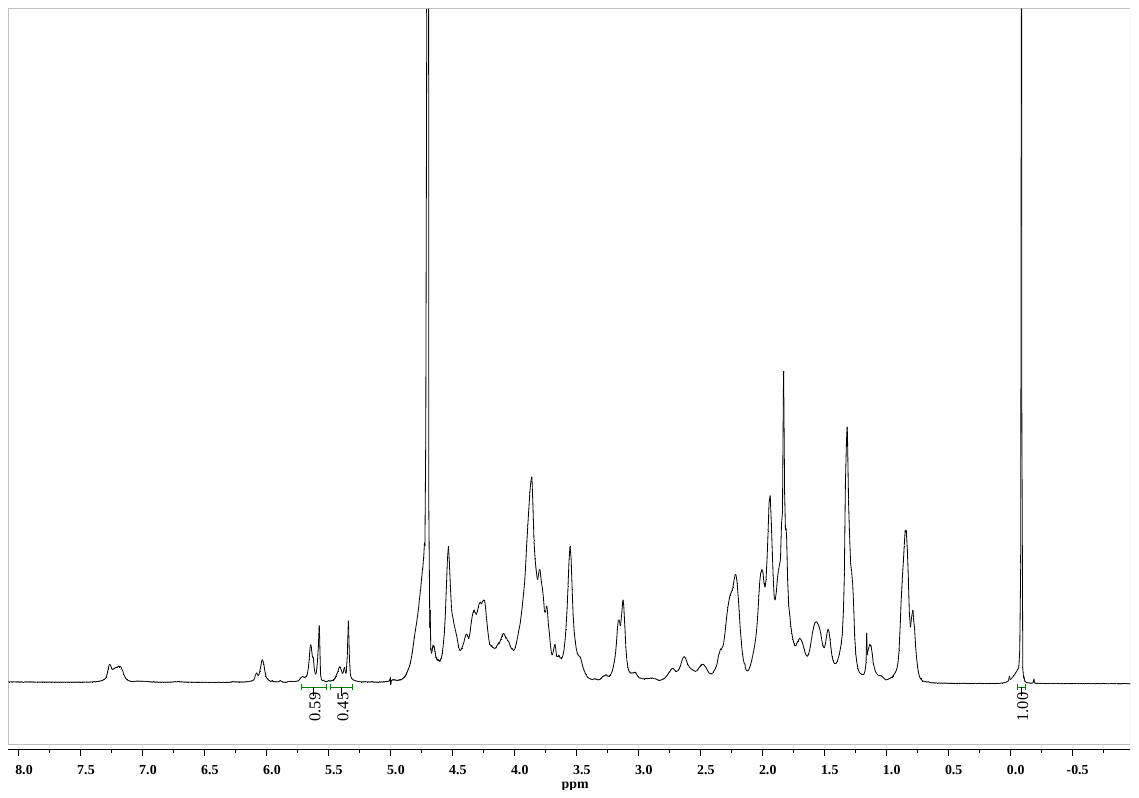

**Fig. S7**. ^1^H NMR spectrum of GelMA in D_2_O.

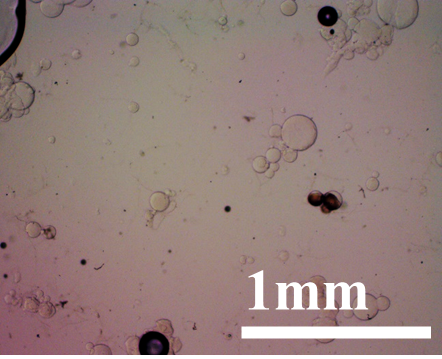

**Fig. S8**. The morphology of the as-synthesized gelatin microspheres.

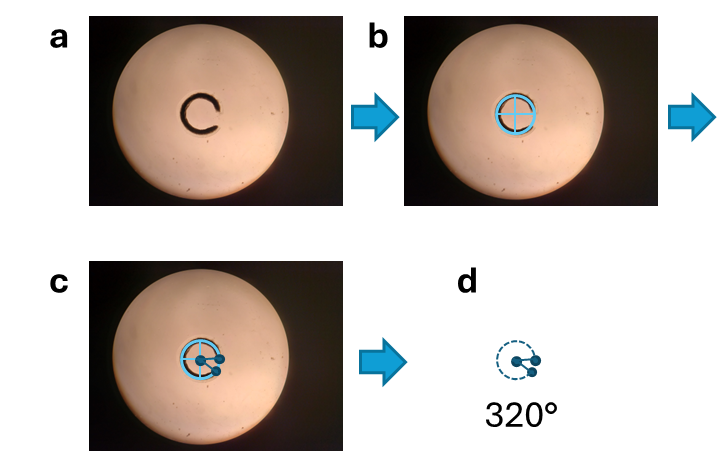

**Fig. S9**. Depiction of bending angle measurement. Here, the Day 5 GM photomicrograph from Fig. 3A is used as an example. (a) First, the image is uploaded into PowerPoint. (b) Then, a circle with crosshairs is fitted to the contours of the printed construct. (c) This overlay is then copied into ImageJ and the angle tool is used to find the angle between the ends of the construct. The middle of the crosshairs is used as the vertex. (d) The bending angle is then measured and recorded.
